# New high-resolution maps show that rubber causes significant deforestation

**DOI:** 10.1101/2022.12.03.518959

**Authors:** Yunxia Wang, Peter M. Hollingsworth, Deli Zhai, Christopher D. West, Jonathan Green, Huafang Chen, Kaspar Hurni, Yufang Su, Eleanor Warren-Thomas, Jianchu Xu, Antje Ahrends

## Abstract

Understanding the impacts of cash crop expansion on natural forest is of fundamental importance. However, for most crops there are no remotely-sensed global maps^1^, and global deforestation impacts are estimated using models and extrapolations. Natural rubber is an example of a major commodity for which deforestation impacts have been highly uncertain, with estimates differing more than five-fold^1–4^. Here we harnessed earth observation satellite data and cloud computing^5^ to produce the first high-resolution maps of rubber and associated deforestation covering all Southeast Asia. Our maps indicate that rubber-related forest loss has been significantly underestimated in policy, by the public and in recent reports^6–8^. Our direct remotely-sensed observations show that deforestation for rubber is two to threefold higher than suggested by figures currently widely used for setting policy^4^. With over 3.76 million hectares of forest loss for rubber since 1993 (2.77 [2.5-3 95% CI] million hectares since 2000), and over 1 million hectares of rubber plantations established in Key Biodiversity Areas, the impacts of rubber on biodiversity and ecosystem services in Southeast Asia are extensive. Thus, rubber deserves more attention in domestic policy, within trade agreements and in incoming due diligence regulations.

---

Around 90-99% of tropical deforestation is linked to the production of global commodities such as beef, soy, oil palm, natural rubber, coffee and cocoa ^9^. Understanding the impacts of individual commodities on natural forests is of fundamental importance for targeted policies and interventions. However, with relatively few exceptions – most notably oil palm and soy^1,10^ – directly observed global or regional maps derived from satellite imagery are unavailable for most commodities. Instead, commodity-specific global deforestation is typically estimated using models^11,12^ and extrapolations^13,14^ with large levels of uncertainty.

Natural rubber is an example of a commodity whose impacts on forests have remained poorly understood despite its economic importance^15^ and the potential for widespread deforestation, land degradation and biodiversity loss^13,16–21^. Natural rubber is used in the manufacture of almost three billion tyres per year^15,22^, and continued and increasing global demand is driving land use conversion in producer countries^14^. Production is primarily located in Southeast Asia (90% of the global production^23^), with the remainder coming from South and Central America and more recently also West and Central Africa^24^. Rubber is produced from the latex of a tropical tree (*Hevea brasiliensis*), and the spectral signature of rubber plantations is similar to forest^25^, making it challenging to identify conversion of natural forest to rubber plantations from space. In addition, around 85% of global natural rubber is produced by smallholders^26^, meaning that the plantations are scattered and often below 5 ha in size, increasing the challenge of detecting them from satellite images and capturing them in national crop statistics. Consequently, the impacts of rubber are surrounded by uncertainty and estimates of rubber-driven deforestation differ by more than five-fold: from less than 1 million ha almost globally between 2005 and 2018^3^ to more than five million ha between 2003 and 2014 in continental South-East Asia alone^2^. Direct observations based on remote sensing have previously only existed for subsets of Southeast Asia^2,27,28^, individual countries^1,29^, or subnational areas^30^, and most are outdated, so do not reflect the current risk.

Currently, the most widely used dataset to estimate global rubber-related deforestation has been derived using a ‘land balance’ model^11^. This model combines remotely sensed data on tree cover loss with non-spatial estimates of crop expansion, derived mainly from national-scale statistics. The ‘land balance’ approach means that tree cover loss is not spatially linked to commodity expansion, and therefore is not a substitute for more accurate products that provide spatially explicit estimates of crop expansion into forest areas, as explicitly acknowledged by the authors^31^. The ‘land balance’-derived data^3,4^ suggest that rubber is a relatively minor problem when compared to the impact of other major forest risk commodities, with palm oil and soy accounting for seven to eight times more deforestation than rubber; and in UK imports^6^ for 20 and 57 times more deforestation, with rubber sitting on a par with nutmeg. This has contributed to the reduced the attention that rubber has received as a driver of deforestation compared to other commodities and has led policy makers to question the need to include rubber in the European Commission’s proposal for a regulation on deforestation-free products^7^, and secondary legislation associated with UK Environment Act Schedule 17 aimed at addressing illegal deforestation. However, given the inherent uncertainty in model-based estimates, there is an urgent need for robust evidence to provide guidance for policy interventions to avoid rubber being prematurely excluded from key policy processes and interventions, particularly as review of policy will not be due for several years.

Here we present up-to-date analyses and provide the first Southeast Asia wide maps of rubber and associated deforestation, encompassing over 90% of the natural rubber production volume. We used the latest high-resolution Sentinel-2 imagery (at a spatial resolution of 10 m) to map the extent of rubber across all Southeast Asia in 2021. Our approach is based on the distinctive phenological signature of rubber plantations which allows them to be distinguished from typical tropical forests based on leaf-fall and regrowth, which occur in specific time windows that differ by region. To tackle the challenge of heavy cloud cover in the region we use multi-year imagery composites. We track the deforestation history of locations occupied by rubber in 2021 using historical Landsat imagery and a spectral-temporal segmentation algorithm (LandTrendr)^32^. Here, we use the term ‘deforestation’, but this can include other types of tree cover loss if that tree cover (e.g., agroforests, plantation forests, agricultural tree crops and rubber itself) was established prior to the 1980s.

## First rubber map for all Southeast Asia

Our results show that mature rubber plantations occupied an area of 14.5 [5.6–23.4 95% CI] million hectares in Southeast Asia in 2021, with over 70% of the production area situated in Indonesia, Thailand and Vietnam. Other significant areas were situated in China, Malaysia, Myanmar, Cambodia, and Laos (Table 1, Fig. 1 A). The rubber maps achieved an overall classification accuracy of 0.91 with a producer’s accuracy of 0.83 (Extended Data Table 1). Our estimates are consistent with the sum of national statistics reported to the Forest and Agriculture Organization of the United Nations (FAO), according to which the total area of harvested rubber in the above eight countries was 10.18 million ha in 2020^23^. Due to the currently low global rubber price many plantations may not be harvested, meaning that although our mean estimate is higher than the values reported to the FAO, there is a broad alignment, and our lower confidence interval is in fact exceedingly conservative. Our estimates are also generally within the bounds estimated by two other recent remote sensing studies for rubber^2,28^ (Table 1).

**Table 1.** Area estimates of rubber plantations for individual countries in Southeast Asia. For China only the main production areas are included (Xishuangbanna and Hainan). 95% confidence intervals were calculated using accuracy estimates presented in Extended Data Table 1. Hurni & Fox (2018) derived both standard mapped figures and error-corrected figures (indicated by an asterisk). For Thailand their figures only include northeast Thailand and for Vietnam only areas south of Hanoi.

| Country | Rubber (ha) | % Rubber | Rubber in KBA (ha) | % Rubber in KBA | FAO 2020 harvested rubber (ha)<sup>23</sup> | Xiao et al. 2021 (rubber in 2018, ha)<sup>28</sup> | Hurni & Fox 2018 (rubber in 2014, ha)<sup>2</sup> |  |
| --- | --- | --- | --- | --- | --- | --- | --- | --- |
| Indonesia | 5,134,276 | 35% | 397,347 | 8% | 3,668,735 | NA | NA | NA |
| Thailand | 3,742,929 | 26% | 291,516 | 8% | 3,292,671 | 4,650,000 | 1,429,487 | 2,861,400* |
| Vietnam | 1,605,900 | 11% | 59,367 | 4% | 728,764 | 740,000 | 912,696 | 1,916,600* |
| China | 1,097,077 | 8% | 58,067 | 5% | 745,000 | NA | NA | NA |
| Malaysia | 985,012 | 7% | 49,380 | 5% | 1,106,861 | NA | NA | NA |
| Myanmar | 779,546 | 5% | 84,561 | 11% | 323,956 | 680,000 | NA | NA |
| Cambodia | 618,039 | 4% | 117,665 | 19% | 310,877 | 200,000 | 917,446 | 2,974,300* |
| Laos | 573,964 | 4% | 49,115 | 9% | NA | 700,000 | 260,471 | 765,600* |
| **Southeast Asia** | **14,536,742 ± 8,930,998 (95% CI)** |  | **1,107,018** | **7.62% ± 4.68% (95% CI)** | **10,176,864** |  |  |  |

**Fig. 1.**
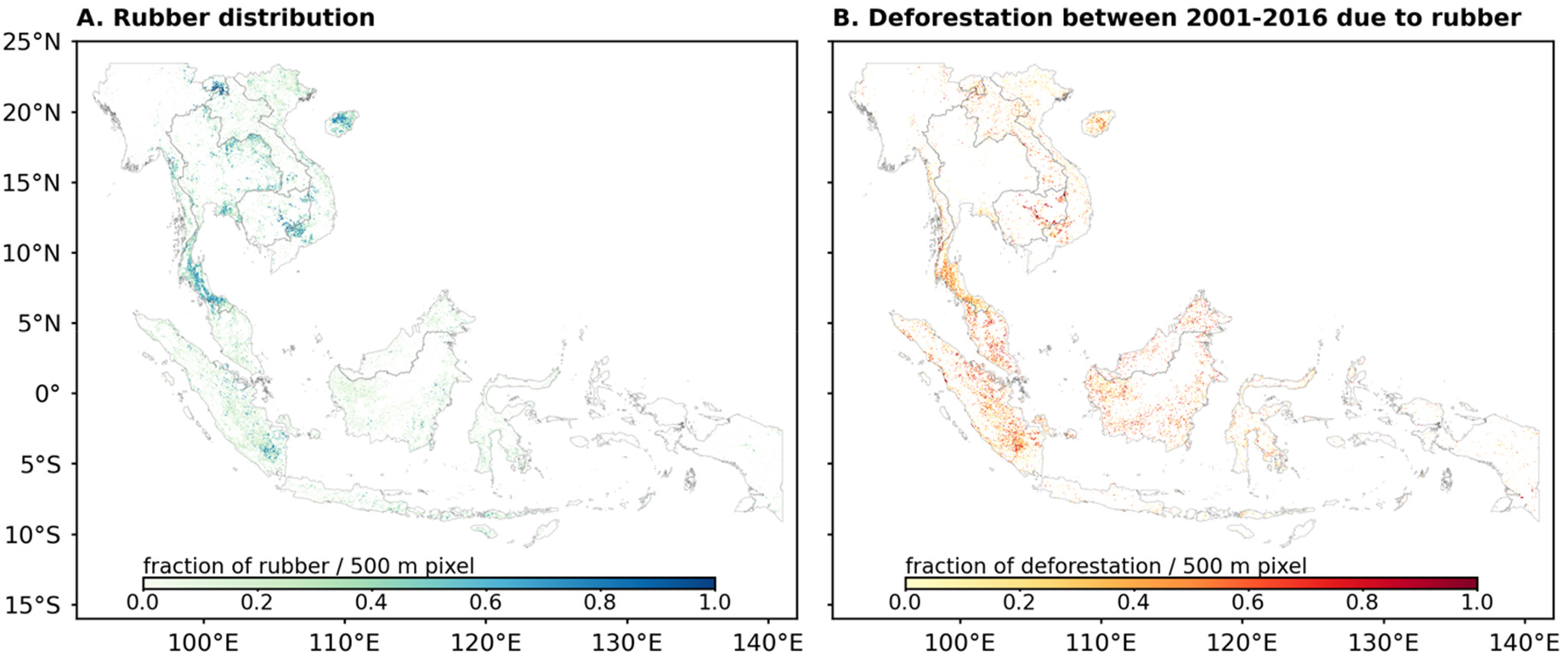
Rubber distribution in 2021 (A) and associated deforestation (B) across Southeast Asia. For better visualization, the rubber map (A) was aggregated to 500 m resolution by calculating the proportion of 10 m rubber pixels in each 500 m pixel; the deforestation due to rubber map (B) was aggregated to 500 m by calculating the proportion of 30 m deforestation pixels within each 500 m pixel. The maps in their original resolution are available at https://wangyxtina.users.earthengine.app/view/rubberdeforestationfig1.

## Significant deforestation due to rubber

We used time-series Landsat imagery to identify the deforestation date for all areas classed as rubber in 2021 in two categories: pre-2000 and 2001-2016 (overall classification accuracy of 0.78; Extended Data Table 2). Specifically, we used the LandTrendr algorithm^33^, which identifies break points in the pixels’ spectral history (Normalized Burn Ratio), indicating a sudden change from forest or other types of tree cover to bare or burnt ground. We only used the first breakpoint, going as far back in time as the imagery allows (1988), meaning that we only include rotational plantation clearance into the deforestation estimate if these plantations were established prior to 1988.

Our data show that rubber led to significant deforestation across all of Southeast Asia (Fig. 1 B). In total, we estimate that 3.76 million ha of forest have been cleared for rubber between 1993 and 2016. Almost three quarters of this forest clearance occurred since 2001 (2.77 [2.53 - 3.01 95% CI] million ha), meaning that around one fifth of the rubber area in 2021 was associated with deforestations occurring after 2000 (Extended Data Table 3). In addition, over 1 million ha of rubber plantations in 2021 were situated in Key Biodiversity Areas^34^, globally important for the conservation of biodiversity (Table 1).

In terms of individual countries, both historically and since 2001, deforestation was highest in Indonesia, followed by Thailand and Malaysia (Figs. 2 and 3). While these three countries account for over two-thirds of the total rubber-related deforestation in Southeast Asia between 2001-2016, significant deforestation also occurred in Cambodia since 2001, where almost 40% of rubber plantations are associated with deforestation (Fig. 2) and 19% with deforestation in Key Biodiversity Areas (Table 1).

**Fig. 2.**
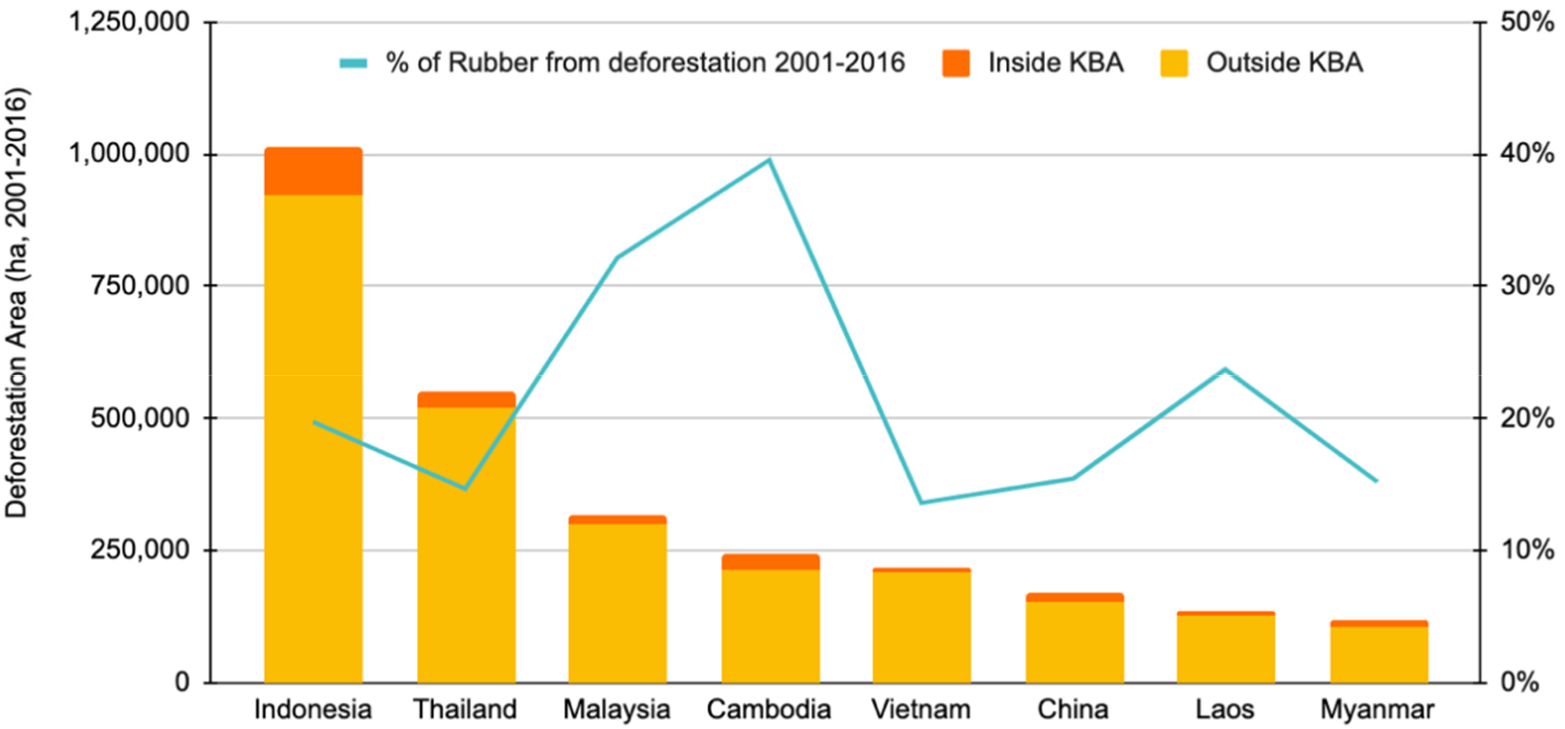
Area of rubber-related deforestation between 2001-2016 for individual countries in Southeast Asia. The bars show the cumulative area of deforestation (2001-2016) for rubber plantations in 2021 and orange areas are the fraction of deforestation that occurred inside Key Biodiversity Areas (KBA)^34^. The blue line shows the percentage of the total national rubber area in 2021 that was associated with deforestation between 2001-2016 (the percentage is given on second y-axis). The figures for China only include its main production areas (Xishuangbanna and Hainan).

**Fig.3.**
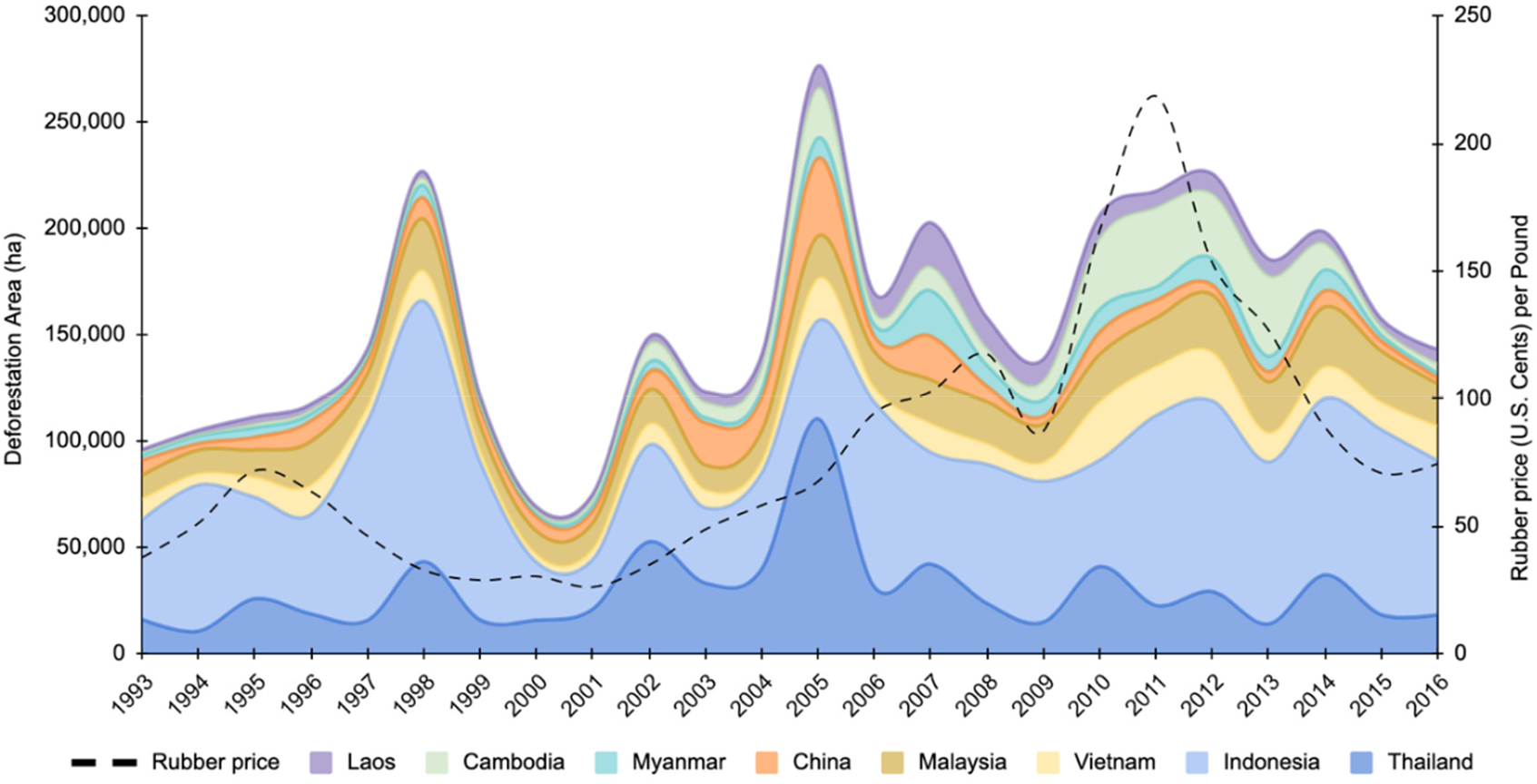
Total area of rubber-related deforestation in Southeast Asia between 1993-2016. The colours show the fraction of overall deforestation that occurred in individual countries. While most deforestation occurred in Indonesia and Thailand and while the deforestation trends are similar across countries, the fraction of deforestation occurring in continental Southeast Asia (mainly Cambodia) has increased over the last decade. Rates of rubber-deforestation in some countries were strongly correlated with the global rubber price (black dashed line, second y axis): simple Pearson’s correlations (i.e., without accounting for a potential temporal lag) were Cambodia R=0.83, Vietnam R=0.75, Malaysia R=0.61, Laos R=0.58, Myanmar R=0.54, and Indonesia R=0.54.

## Rubber deforestation is underestimated

Recent estimates of deforestation embedded in rubber, intended for inform policy by the EU^7^, G7^8^ and UK^6^, all used the data generated by Pendrill et al. (2019)^11^, which place total rubber-related deforestation (in 135 countries, including all major rubber producers except China and Laos) between 2005-2017 at below 700,000 ha. Translating to an average annual deforestation of 53,000 ha (Table 2), these estimates lie several-fold below the estimates of this and other studies based on spatially explicit data - in the case of Cambodia several hundred-fold (Table 2). An update of the Pendrill et al. data^3^ now provides an almost 30-fold higher estimate for deforestation in Cambodia (Table 2), but still places total quasi-global rubber-related deforestation between 2005-2018 below 1 million ha. In contrast, the World Resources Institute^1^ estimated that rubber replaced 2.1 million ha of forest 2001-2015 in just seven countries, which account for less than half of the global natural rubber production, and Hurni and Fox (2018)^2^ estimated that rubber replaced more than 5 million ha of forest in continental Southeast Asia alone. Although our estimates are in fact conservative compared to these other estimates, and although none of the figures can be directly compared as they refer to somewhat different time periods and different definitions of forest, it is of critical note that even our lower 95% confidence interval still greatly exceeds (three-fold) the model-based estimates currently widely used to guide policy and to calculate deforestation footprints. Furthermore, even if we replaced our estimates for Indonesia and Malaysia with those of Pendrill et al. (2019), the two countries in which Pendrill et al. (2019) attempted to exclude plantation rotation from deforestation totals, our annual rubber-deforestation totals would still be more than twice as high (Extended Data Note).

**Table 2.** Comparison of rubber-related deforestation estimates generated by this and other studies. The dataset in grey (first row) has been used to guide deforestation policy^7^ and to calculate individual countries’ imported deforestation^6,8^. Hurni & Fox (2018) derived both standard mapped figures and error-corrected figures (indicated by an asterisk).

|  | Method | Definition of 'forest' | Time period | Reference area | Rubber related deforestation in 1000 ha yr<sup>-1</sup> |  |  |  |  |
| --- | --- | --- | --- | --- | --- | --- | --- | --- | --- |
|  |  |  |  |  | Total in reference area | Indonesia | Thailand | Malaysia | Cambodia |
| Pendrill et al. 2020<sup>4</sup> | Land-balance model | Tree cover >=25% (Hansen et al. 2013) | 2005-2017 | 135 tropical countries, incl. all major rubber producers (except China and Laos) | 53 | 22 | 9 | 5 | 0.1 |
| Pendrill et al. 2022<sup>3</sup> |  |  | 2005-2018 |  | 52 | 23 | 6 | 5 | 3 |
| Goldman et al. 2020<sup>1</sup> | Mix of spatially explicit data | Tree cover >=30% (Hansen et al. 2013) | 2001-2015 | Brazil, Cambodia, Cameroon, Democratic Republic of the Congo, India, Indonesia, and Malaysia | 140 | 64 | NA | 48 | 22 |
| Hurni & Fox 2018<sup>2</sup> | Remote sensing | Internal classifier | 2003-2014 | Mainland Southeast Asia | 135 | NA | NA | NA | 69 |
|  |  |  |  |  | 437* |  |  |  | 232* |
| Grogan et al. 2019<sup>29</sup> | Remote sensing | Tree cover >=10% (Sexton et al. 2013) | 2001-2015 | Cambodia |  | NA | NA | NA | 34 |
| This study | Remote sensing | ESA WorldCover 10-m 2020 v100 (tree cover >=10%) | 2001-2016 | Southeast Asia | 173 ± 15 | 63 | 34 | 20 | 15 |

## Discussion

Here we provide the first high-resolution maps for rubber and associated deforestation between 1993-2016 for all Southeast Asia. We show that rubber has led to several million ha of deforestation, and that the global data^3,4^ currently widely used in setting deforestation policies are likely to severely underestimate the scale of the problem. Whilst very helpful for providing a holistic assessment of the role of agricultural commodities in driving tropical and subtropical deforestation across the globe, these and other model-based data are not a substitute for spatially explicit estimates of crop expansion into natural forests^31^. Our estimates lie several-fold above these data despite only covering Southeast Asia and not, for example, West and Central Africa, where there has been significant recent rubber expansion, likely driving deforestation^24^.

Due to the heterogenous data landscape with greatly variable accuracy across crops, the impacts of crops on deforestation cannot be reliably compared. The findings of this study would place rubber deforestation above the impacts found for coffee, and, contrary to previously assumed, above the impacts of cocoa^1,4^. While still lower than the impacts of oil palm, not so by a factor of 8-10 as has been previously suggested^1,4^ and instead only by a factor of 2.5-4 (also noting that here we are comparing our data for Southeast Asia only with global estimates for these other crops). However, these comparisons are difficult to make, not least because the estimated impacts of cocoa also differ threefold between studies^1,4^ with cocoa being another example of a crop for which there are no global remotely-sensed maps.

Our figures are likely to be conservative: First, we used 2021 as the reference year and hence do not capture deforestation for rubber if by 2021 the rubber plantation was converted to a different land use. Since there was a rubber price boom in the first decade of this millennium, followed by a price crash since 2011^35^, it is possible that in the meantime some rubber area has been converted to other more lucrative land uses^36^, which will not be included in our estimates. Second, we used the ESA global tree cover map^37^ as a mask for mapping rubber plantations. If rubber areas were not picked up as tree cover by this map, they are also excluded from our estimates. Third, due to continuous cloud cover small areas in the region lacked clear Sentintel-2 images and had to be excluded (especially in Indonesia). While the area that had to be excluded due to cloud cover was very small, a noteworthy wider issue is that our maps are potentially more accurate for mainland Southeast Asia than for insular Southeast Asia, where in addition to more persistent cloud cover, other challenges were present in the form of a less predictable rubber phenology and complex land use pattens and trajectories. Finally, we only map mature rubber; younger rubber plantations (around <5 years old) are excluded.

We have considered and accommodated possible areas of ambiguity that might otherwise lead to an overestimation of deforestation using our method. First, rotational plantation and tree crop clearing and replanting may erroneously be classed as deforestation. This is a key issue, which is notoriously difficult to address and hence also affects other studies^1,11^ (Extended Data Note). The issue is likely to be particularly important in Indonesia, Malaysia, and Thailand where rubber and other plantations have a longer history of planting. To address this, we only use the first deforestation date and ignore subsequent pixel changes, meaning that this problem would only apply to plantations and tree crops established prior to, and mature by, 1988. This baseline is relatively conservative. Second, deforestation may have occurred for a different land use, with the area then subsequently being converted to rubber. This may indeed affect our data, but the issue will be smaller for rubber than for example for oil palm, which boomed and expanded more recently^38^, possibly replacing other land uses in addition to forests, whereas rubber is a crop with a longer history in the area and a greater plantation longevity of c. 25 years^30^. Third, the vegetation in some pixels may have undergone some type of disturbance in the rubber defoliation time window, followed by regrowth in the rubber refoliation window, leading to them having the characteristic phenology signature of rubber and erroneously being classed as such. To exclude such pixels and increase the accuracy of our analysis we created a ‘disturbance’ mask (see Methods). Thus overall, we consider that the estimates of deforestation due to rubber plantations that we have provided are more likely to be an underestimate than an overestimate of the scale of the issue.

The current estimates for deforestation caused by rubber^3,4^ used for policy considerations in the EU^7^ and UK^6^ are based on a land-balance model^11,12^. Such models typically allocate total deforestation area to different commodities based on national (or sub-national, e.g. in the case of this model for Brazil and Indonesia) reports of crop expansion^11^. This can lead to significant over or underestimates of the role of different crops in driving deforestation^31^. First, crop expansion statistics are hampered by uncertainties and inconsistent reporting across crops and countries. Secondly, while the total area of a crop can remain stable, its actual place of occupancy may change^31^. This is highly relevant to rubber as oil palm has expanded into traditional rubber growing areas^39^, with new compensatory rubber plantations being established elsewhere, e.g., in uplands^18,30^ and often climatically marginal areas^16^, where they may be associated with deforestation. Thus, while the use of extrapolation^13,14^ and model-based^11,12^ approaches provide some form of estimation for the extent of deforestation due to rubber plantations, we advocate caution in their interpretation. Instead, where available, we argue for the use of results from direct observations of the dynamics of crop production systems (e.g., using remotely-sensed satellite imagery), thereby greatly increasing the accuracy of deforestation estimates.

In terms of future projections of the impact of rubber and the time critical need for deforestation legislation, it is likely that demand for natural rubber will continue to increase^15^. Synthetic alternatives or other natural sources are not a perfect substitute^40,41^, and, being based on petrochemicals primarily derived from crude oil, they are also considered more environmentally harmful. Natural rubber on the other hand is a renewable resource with the potential to contribute to climate change mitigation^42^ and to benefit the livelihoods of smallholder farmers^43^. However, if not regulated carefully, rubber can have severe negative consequences for both the environment^13,16–21^ and livelihoods^26,44^. Our deforestation data also suggest that the assumed ‘breathing space’^36^ generated by the currently low rubber price may be false, with continued (and volatile) deforestation for rubber since 2011, a problem that is likely to increase when rubber prices rise again.

Given the significant rubber related deforestation demonstrated here, it is encouraging that an increasing number of global initiatives aim to address this. A frequently voiced concern by critics is that it is very difficult for rubber operators to trace their supply chains and that any deforestation regulations would present a disproportionate burden for rubber operators. Contrary to for example oil palm where there is a limited time window (c. 24 hours) between harvest and mill processing, the raw rubber harvest has greater longevity and hence can travel several hundred km and change hands between half a dozen or more aggregators before arriving at processing facilities^45^. Another critically important point is the need to ensure that the poorest countries and importantly smallholders are not disadvantaged by deforestation regulations, as contrary to larger companies they may not be able to afford the premiums for certified sustainable production. While this applies to all commodities, it is a particularly important consideration for commodities that are strongly linked to smallholder livelihoods and development prospects, such as rubber. Recent initiatives for example by the Forest Stewardship Council have demonstrated that these challenges can be overcome when farmers are organised in groups, with an additional benefit being that farmer cooperatives can negotiate a joint price to buffer their livelihoods against the volatile global rubber price. In addition, whilst supply chains are indeed complex and challenging to trace, the high-end rubber processing side is dominated by very few and identifiable actors. Around 70% of the global natural rubber production is used in tyres with a few major tyre companies accounting for the majority of global consumption^15^. Many of these are already part of the Global Platform for Sustainable Natural Rubber (GPSNR) – an international multistakeholder membership organisation committing to lead improvements in socioeconomic and environmental performance of the natural rubber value chain. Further work would be needed to make connections between rubber-driven deforestation and specific EU supply chains, but in the absence of such information it should be assumed that the EU is significantly exposed to rubber-deforestation, with over 40% of EU natural rubber imports coming from Indonesia, with much of the remainder coming from Thailand and Malaysia^46^, i.e. countries that according to our data experience some of the most significant rubber-driven deforestation. In addition, the lack of traceability information at the current time provides a further argument for the inclusion of rubber in regulatory processes in order to improve traceability and to provide an opportunity for the EU supply chain to support sustainable production.

In summary, we believe that rubber merits more consideration in policies and processes that aim to reduce commodity driven deforestation, and that it is vitally important to use the best available evidence on the scale of the problem. The issue outlined here for rubber is of fundamental importance in its own right because rubber is responsible for millions of hectares of deforestation. However, we also highlight the wider need to enhance the evidence base available to inform policy decisions. There is an opportunity for increased clarity and rigorous quantification in the extent of environmental degradation caused by major cash crops that is increasingly possible using remotely sensed earth observation.

## Methods

We used Sentinel-2 imagery to produce an up-to-date distribution map of rubber plantations for all Southeast Asia and mapped this against time-series data from Landsat images between 1988-2021 to identify the historical deforestation date for areas of rubber in 2021.

### Sentinel-2 imagery

Sentinel-2 is an optical multispectral imaging mission from the Copernicus Program headed by European Commission in partnership with European Space Agency (ESA). It acquires very high-resolution multispectral imagery with a global revisit frequency of 5 days. In this study, we used the Sentinel-2 level-2A Surface Reflectance (SR) imagery^1^ obtained through Google Earth Engine^2^ to map the extent of rubber plantations in Southeast Asia. Sentinel-2 SR imagery has been corrected for atmospheric influences with the ‘Sen2Cor’ processor algorithm^3–5^. To remove clouds and cloud shadows, we applied the ‘QA60’ cloud mask band from the Sentinel-2A SR imagery, and Sentinel-2 Cloud Probability datasets^4^ where pixels with cloud probability greater than 50% are considered as clouds. Cloud shadows are defined as areas of cloud projection intersection with low-reflectance near-infrared pixels. The full details of masking clouds and cloud shadows can be found at https://developers.google.com/earth-engine/tutorials/community/sentinel-2-s2cloudless.

Sentinel-2 SR images acquired between 2020-2022 were used as inputs for mapping rubber extent. For each image, we selected ten spectral bands and computed seven vegetation indices. The selected bands included four 10 meter resolution bands (Blue: B2, Green: B3, Red: B4 and Near-Infrared: B8) and six 20 meter resolution bands (Red Edge bands^6^: B5, B6, B7, B8A, Short-wave Infrared bands: B11, B12). The seven vegetation indices included Normalized Difference Vegetation Index (NDVI), Normalized Difference Water Index (NDWI), Renormalization of Vegetation Moisture Index (RVMI), Normalized Burn Ratio (NBR), Modified Normalized Burn Ratio (MNBR), Soil Adjusted Vegetation Index (SAVI) and Enhanced Vegetation Index (EVI). All the spectral bands and vegetation indices were resampled to 10 m resolution for further analysis. The equations used for calculating the vegetation indices are as follows:

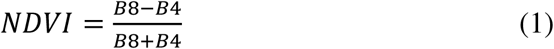

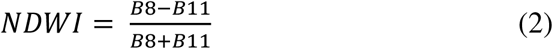

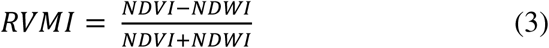

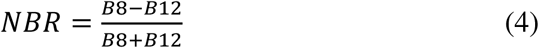

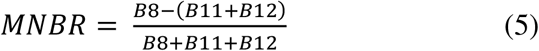

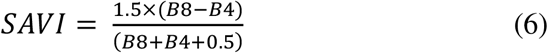

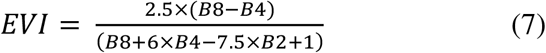

### Mapping the extent of rubber plantations

We designed a novel phenology-based methodology to map rubber plantations across Southeast Asia. Unlike tropical rainforests and other tree plantations, rubber plantations shed their leaves during the drier and colder season and subsequently regain their leaves. For instance, in mainland Southeast Asia defoliation generally occurs during January-February and the subsequent refoliation during March-April. In Indonesia the occurrence of the dry season is more spatially heterogenous^7^. In some areas the lowest monthly precipitation occurs during June-September. Backed by sample data^8,9^ we made the assumption that in these areas the defoliation occurs during June-September with the subsequent refoliation occurring during October-December (Extended Data Fig.2). We refer to the areas where rubber defoliation occurs during January-February as region-A, and where rubber defoliation occurs in June-September as region-B.

The unique phenology of rubber gives it distinct spectral characteristics, making it distinguishable from other tree cover using satellite imagery. In this study, we used a tree cover mask from the ESA global land cover map^10^ (The European Space Agency WorldCover 10 m 2020 product) and classified tree cover into rubber and other tree cover based on the spectral differences between rubber refoliation and defoliation stages (using Sentinel-2 imagery). For the defoliation stage, we generated a composite image using 15% NDVI percentile of all images acquired during January-February in 2021 and 2022 for region-A, and during June-September in 2020 and 2021 for region-B. For the refoliation stage, we used the 85% NDVI percentile composite of all images acquired during March-April in 2021 and 2022 for region-A, and during October-December in 2020 and 2021 for region-B. Each composite image contained 17 variables, including 10 spectral bands and 7 vegetation indices (see Sentinel-2 imagery above). The composite image difference between the refoliation and defoliation stages was subsequently used as input for a Random Forest machine learning classification.

To run the machine learning classification, individual samples are required for each classification category. In this study, we collected a total of 2,010 rubber points and 1,816 evergreen forest points from ground-truthed points, publications^11,12^ and random sampling points, which we visually interpreted using high-resolution satellite imagery through the software Collect Earth Online^13–15^ (CEO). For the latter, we randomly sampled 1,000 forest points and 1,000 rubber points from ground-truth data collected in 2010 (World Agroforestry Centre Southeast Asia) and visually interpretated and subsequently re-labelled these points for the year 2021 through CEO and Google Earth^16^. In CEO, the visual interpretation was based on the Mapbox Satellite imagery base map, 2021 monthly Planet NICFI images^17^ (Norway’s International Climate and Forests Initiative satellite data program) and yearly composite images for January-February and March-April from Sentinel-2 (2017-2021)^1^ and Landsat-5-7-8 (1988-2016)^18^. First, we assigned each sample point to a land cover class for the year 2021.

If the land cover was rubber, we further identified the deforestation date for that point using historical Landsat images^18^, starting from 1988. Where available, additional very high-resolution imagery from Google Earth^16^ was used to facilitate the interpretation process.

We randomly split all sampling points into 80% as training samples for mapping rubber, using the remaining 20% to validate the final rubber map.

Disturbances such as degradation or plantation removal could potentially produce similar spectral features to rubber phenology, leading to commission errors. To reduce commission errors, we removed all rubber pixels where this may have occurred using a 2021 primary forest mask and a no-disturbance mask (Extended Data Fig.1). The 2021 primary forest mask was created by using the 2001 primary forest layer from Turubanova, et al. (2018)^19^ and removing areas of subsequent forest loss between 2000-2021 (Hansen Global Forest Change v1.9)^20^. The no-disturbance mask was generated with the following steps: (1) Calculate the NBR index (Normalized Burn Ratio, above equation-4) for all Sentinel-2 images between 2019-2021; (2) Create NBR three-year median composites for March-June, July-September and October-December (region-A) or January-May and October-December (region-B) (yielding three composites for region-A, and two composites for region-B); (3) Extract the values of NBR composites for all the rubber samples; (4) Plot the NBR values and calculate the 5% percentile thresholds for individual composites, meaning 95% of rubber samples’ NBR values are above these thresholds; (5) Apply the thresholds to all three (region-A) or two (region-B) NBR composite images, resulting five binary images (1: no disturbance, 0: potential disturbance). If a pixel was classed as 1 in all three (region-A) or two (region-B) binary images, it was considered as not disturbed. A 5 by 5-pixel majority filter was applied to the no-disturbance mask to remove isolated pixels.

In summary, we developed a novel approach, which involves classifying an ESA tree cover baseline map^10^ into rubber and other tree cover based on phenology, and removing any pixels that are potentially confounded by disturbance using a primary forest mask and a no-disturbance mask, which we generated specifically for this purpose. We also applied a post-classification 5 by 5-pixel majority filter to the resulting map, and a minimum patch size threshold of 0.5 ha to reduce pixel-level classification noise and to remove classification artifacts.

### Identifying the deforestation date

We tracked the first historical deforestation date for all rubber plantations mapped in 2021. This was done using the LandTrendr spectral-temporal segmentation algorithm^21,22^ (a Landsat-based algorithm for the detection of trends in disturbance and recovery). LandTrendr characterises the history of a Landsat pixel by decomposing the time series into a series of bounded line segments (i.e. trends over several years) and identifying the break points between them. These linear segments and breakpoints allow for the detection the greatest pixel-level change (e.g. deforestation) and therewith for the identification of the year in which this greatest spectral change occurred.

In this study, we ran LandTrendr GEE API^22^ (a JavaScript module developed in Google Earth Engine, https://emapr.github.io/LT-GEE/api.html) using the annual time-series index from USGS Landsat Surface Reflectance Tier 1 datasets. The clouds and cloud shadows were masked using the CFMASK^23^. A medoid approach was used to generate the annual composite image. This approach uses the value of a given band that is numerically closest to the median of all the available images for each year. In this study, we used the time-series NBR index (Normalized Burn Ratio, equation-4 above) from 1988 to 2021 for the temporal segmentation. The deforestation date was identified as the end year of the linear segment with the largest slope (greatest loss). As an additional constraint, we imposed a minimum start NBR value for this linear segment of over 0.595, thereby reducing the risk of including the clearance of old plantations (planted before 1988) as deforestation. Any deforestation pixels below this threshold were excluded from our deforestation estimates. We tested a range of NBR thresholds and selected this one as it provided maximum overall accuracy. We also excluded pixels with a deforestation date later than 2016 because it takes around 5 years for rubber plantations to be identifiable from the satellite imagery following deforestation.

We validated the deforestation date map using the deforestation dates of rubber points collected through Collect Earth Online^15^ (see section above on mapping the extent of rubber plantations). In total, there were 704 rubber points with deforestation dates, 80% of which were from deforestation before 1990. As we did not have ground-truthed deforestation points for all years, we grouped the deforestation dates into two broader time periods (pre-2000 and between 2001-2016).

### Deforestation in Key Biodiversity Areas associated with rubber

We further explored the potential impacts of rubber and associated deforestation on regional biodiversity. To do this, we clipped our maps of rubber and associated deforestation to a shapefile for Key Biodiversity Areas (KBA)^24^, and then calculated the area of rubber and associated deforestation within these areas. Key Biodiversity Areas are some of the most critical sites for the conservation of species and habitats globally. Rubber and deforestation in these areas thus poses a significant threat to global biodiversity.

## Data availability

The earth observation datasets that supported the findings of this study are publicly available (e.g., Google Earth Engine data catalogue). The final maps of rubber and associated deforestation will be made publicly available.

## Code availability

The code used for this study will be made available publicly available.

## Acknowledgements

This work was funded by UK Research and Innovation’s Global Challenges Research Fund (UKRI GCRF) through the Trade, Development and the Environment Hub project (project number ES/S008160/1). E.W-T. was supported Natural Environment Research Council NERC-IIASA Collaborative Fellowship (NE/T009306/1). H.C., Y.S. and J.X. were supported by Key Research Program of Frontier Sciences, CAS, Grant No. QYZDY-SSW-SMC014. The Royal Botanic Garden Edinburgh is supported by the Scottish Government’s Rural and Environment Science and Analytical Services Division. We thank C. Ryan, C. Ellis, S. Glaser and N.D. Burgess for comments on the manuscript.

## Author contributions

Y.W. and A.A. conceived the study. Y.W. performed the data analysis with support from D.Z., A.A., C.D.W. and J.G., with D.Z., H.C., K.H, Y.S., E.W-T. and J.X.. contributing ground-truth data. Y.W., A.A. and P.M.H wrote the manuscript. All authors discussed the results and commented on the manuscript.

## Additional information

Supplementary Information is available for this paper.

Correspondence and requests for materials should be addressed to Y.W. or A.A.

## Extended Data legends

Extended Data Note | Definitions of ‘forest’ and ‘deforestation’

Extended Data Table 1 | Confusion matrix for mapping rubber across Southeast Asia.

Extended Data Table 2 | Confusion matrix for mapping deforestation associated with rubber across Southeast Asia.

Extended Data Table 3 | Area of rubber-related deforestation for individual countries in Southeast Asia. The 95% confidence Interval (CI) was calculated using the accuracy estimates presented in Extended Data Table 2.

Extended Data Fig.1 | Methodology flow for mapping rubber (blue), generating non-disturbance mask (green) and estimating deforestation (orange). Different image composites were used for region-A (defoliation between January-February) and region-B (defoliation between June-September). All processing was done in Google Earth Engine.

Extended Data Fig.2 | Rubber phenology regions, grids, and sampling points. As rubber phenology varies across Southeast Asia we divided the study area into two regions using OpenLandMap Monthly Precipitation^7^. Region-A: rubber defoliation was assumed to occur between January-February and refoliation between March-April. Region-B: rubber defoliation was assumed to occur between June-September and refoliation between October-December. The algorithm was run separately for 3 by 3-degree grid cells (in blue). The forest and rubber sample ground-truth points were used for training the algorithm (80%) and subsequently validating the map (20%).

## References

1 Goldman, E., Weisse, M. J., Harris, N. & Schenider, M. Estimating the Role of Seven Commodities in Agriculture-Linked Deforestation: Oil Palm, Soy, Cattle, Wood Fiber, Cocoa, Coffee, and Rubber. (Technical Note. Washington, DC: World Resources Institute., 2020).

2 Hurni, K. & Fox, J. The expansion of tree-based boom crops in mainland Southeast Asia: 2001 to 2014. Journal of Land Use Science 13, 198–219, doi:10.1080/1747423x.2018.1499830 (2018).

3 Pendrill, F., Persson, U., Kastner, T. & Wood, R. (Zenodo, 2022). Doi:10.5281/zenodo.5886600

4 Pendrill, F., Persson, U. & Kastner, T. (Zenodo, 2020). Doi:10.5281/zenodo.4250532

5 Gorelick, N. et al. Google Earth Engine: Planetary-scale geospatial analysis for everyone. Remote Sensing of Environment 202, 18–27, doi:10.1016/j.rse.2017.06.031 (2017).

6 Molotoks, A. & West, C. Which forest-risk commodities imported to the UK have the highest overseas impacts? A rapid evidence synthesis. Emerald Open Research 3, doi:10.35241/emeraldopenres.14306.1 (2021).

7 European Commission. Impact Assessment - minimising the risk of deforestation and forest degradation associated with products placed on the EU market. Part 1/2. https://ec.europa.eu/environment/forests/pdf/SWD_2021_326_1_EN_Deforestation%20impact_assessment_part1.pdf. (Brussels, 2021).

8 The Food and Land Use Coalition. Assessing the G7s international deforestation footprint and measures to tackle it. https://www.foodandlandusecoalition.org/wp-content/uploads/2022/09/Assessing-the-G7s-international-deforestation-footprint-and-measures-to-tackle-it.pdf. (2022).

9 Pendrill, F. et al. Disentangling the numbers behind agriculture-driven tropical deforestation. Science 377, eabm9267, doi:10.1126/science.abm9267 (2022).

10 Descals, A. et al. High-resolution global map of smallholder and industrial closed-canopy oil palm plantations. Earth System Science Data 13, 1211–1231, doi:10.5194/essd-13-1211-2021 (2021).

11 Pendrill, F., Persson, U. M., Godar, J. & Kastner, T. Deforestation displaced: trade in forest-risk commodities and the prospects for a global forest transition. Environmental Research Letters 14, doi:10.1088/1748-9326/ab0d41 (2019).

12 Pendrill, F. et al. Agricultural and forestry trade drives large share of tropical deforestation emissions. Global Environmental Change 56, 1–10, doi:10.1016/j.gloenvcha.2019.03.002 (2019).

13 Warren-Thomas, E., Dolman, P. M. & Edwards, D. P. Increasing Demand for Natural Rubber Necessitates a Robust Sustainability Initiative to Mitigate Impacts on Tropical Biodiversity. Conservation Letters 8, 230–241, doi:10.1111/conl.12170 (2015).

14 Warren-Thomas, E., Ahrends, A., Wang, Y., Wang, M. M. H. & Jones, J. P. G. Rubber needs to be included in deforestation-free commodity legislation. doi:10.1101/2022.10.14.510134 (2022).

15 Laroche, P., Schulp, C. J. E., Kastner, T. & Verburg, P. H. Assessing the contribution of mobility in the European Union to rubber expansion. Ambio 51, 770–783, doi:10.1007/s13280-021-01579-x (2022).

16 Ahrends, A. et al. Current trends of rubber plantation expansion may threaten biodiversity and livelihoods. Global Environmental Change 34, 48–58, doi:10.1016/j.gloenvcha.2015.06.002 (2015).

17 Li, H., Aide, T. M., Ma, Y., Liu, W. & Cao, M. Demand for rubber is causing the loss of high diversity rain forest in SW China. Biodiversity and Conservation 16, 1731–1745, doi:10.1007/s10531-006-9052-7 (2006).

18 Feng, Y. et al. Upward expansion and acceleration of forest clearance in the mountains of Southeast Asia. Nature Sustainability, doi:10.1038/s41893-021-00738-y (2021).

19 Guardiola-Claramonte, M. et al. Hydrologic effects of the expansion of rubber (Hevea brasiliensis) in a tropical catchment. Ecohydrology 3, 306–314, doi:10.1002/eco.110 (2010).

20 Zeng, Z. et al. Highland cropland expansion and forest loss in Southeast Asia in the twenty-first century. Nature Geoscience 11, 556–562, doi:10.1038/s41561-018-0166-9 (2018).

21 Kenney-Lazar, M., Wong, G., Baral, H. & Russell, A. J. M. Greening rubber? Political ecologies of plantation sustainability in Laos and Myanmar. Geoforum 92, 96–105, doi:10.1016/j.geoforum.2018.03.008 (2018).

22 Grammelis, P., Margaritis, N., Dallas, P., Rakopoulos, D. & Mavrias, G. A Review on Management of End of Life Tires (ELTs) and Alternative Uses of Textile Fibers. Energies 14, doi:10.3390/en14030571 (2021).

23 FAO. FAOSTAT Statistical Database, <https://www.fao.org/faostat/en/#data> (2022).

24 Feintrenie, L. Agro-industrial plantations in Central Africa, risks and opportunities. Biodiversity and Conservation 23, 1577–1589, doi:10.1007/s10531-014-0687-5 (2014).

25 Li, Z. & Fox, J. M. Mapping rubber tree growth in mainland Southeast Asia using time-series MODIS 250 m NDVI and statistical data. Applied Geography 32, 420–432, doi:10.1016/j.apgeog.2011.06.018 (2012).

26 Fox, J. & Castella, J.-C. Expansion of rubber (Hevea brasiliensis) in Mainland Southeast Asia: what are the prospects for smallholders? Journal of Peasant Studies 40, 155–170, doi:10.1080/03066150.2012.750605 (2013).

27 Hurni, K., Schneider, A., Heinimann, A., Nong, D. & Fox, J. Mapping the Expansion of Boom Crops in Mainland Southeast Asia Using Dense Time Stacks of Landsat Data. Remote Sensing 9, doi:10.3390/rs9040320 (2017).

28 Xiao, C. et al. Latest 30-m map of mature rubber plantations in Mainland Southeast Asia and Yunnan province of China: Spatial patterns and geographical characteristics. Progress in Physical Geography: Earth and Environment, doi:10.1177/0309133320983746 (2021).

29 Grogan, K., Pflugmacher, D., Hostert, P., Mertz, O. & Fensholt, R. Unravelling the link between global rubber price and tropical deforestation in Cambodia. Nat Plants 5, 47–53, doi:10.1038/s41477-018-0325-4 (2019).

30 Chen, H. et al. Pushing the Limits: The Pattern and Dynamics of Rubber Monoculture Expansion in Xishuangbanna, SW China. PLoS One 11, e0150062, doi:10.1371/journal.pone.0150062 (2016).

31 Persson, M., Kastner, T. & Pendrill, F. Flawed numbers underpin recommendations to exclude commodities from EU deforestation legislation. http://www.focali.se/filer/Focali%20brief_2021_02_Flawed%20numbers%20underpin%20recommendations%20to%20exclude%20commodities%20from%20EU%20deforestation%20legislation.pdf (2021).

32 Kennedy, R. et al. Implementation of the LandTrendr Algorithm on Google Earth Engine. Remote Sensing 10, doi:10.3390/rs10050691 (2018).

33 Grogan, K., Pflugmacher, D., Hostert, P., Kennedy, R. & Fensholt, R. Cross-border forest disturbance and the role of natural rubber in mainland Southeast Asia using annual Landsat time series. Remote Sensing of Environment 169, 438–453, doi:10.1016/j.rse.2015.03.001 (2015).

34 BirdLife International. World Database of Key Biodiversity Areas. Developed by the KBA Partnership: BirdLife International, International Union for the Conservation of Nature, American Bird Conservancy, Amphibian Survival Alliance, Conservation International, Critical Ecosystem Partnership Fund, Global Environment Facility, Re:wild, NatureServe, Rainforest Trust, Royal Society for the Protection of Birds, Wildlife Conservation Society and World Wildlife Fund. March 2022 version. Available at http://keybiodiversityareas.org/kba-data/request. (2022).

35 Su, C.-W., Liu, L., Tao, R. & Lobonţ, O.-R. Do natural rubber price bubbles occur? Agricultural Economics (Zemědělská ekonomika) 65, 67–73, doi:10.17221/151/2018-agricecon (2019).

36 Zhang, J.-Q., Corlett, R. T. & Zhai, D. After the rubber boom: good news and bad news for biodiversity in Xishuangbanna, Yunnan, China. Regional Environmental Change 19, 1713–1724, doi:10.1007/s10113-019-01509-4 (2019).

37 Zanaga, D., Van De Kerchove, R., De Keersmaecker, W., Souverijns, N., Brockmann, C., Quast, R., Wevers, J., Grosu, A., Paccini, A., Vergnaud, S., Cartus, O., Santoro, M., Fritz, S., Georgieva, I., Lesiv, M., Carter, S., Herold, M., Li, Linlin, Tsendbazar, N.E., Ramoino, F., Arino, O. ESA WorldCover 10 m 2020 v100. (doi:10.5281/zenodo.5571936). (2021).

38 Meijaard, E. & Sheil, D. The Moral Minefield of Ethical Oil Palm and Sustainable Development. Frontiers in Forests and Global Change 2, doi:10.3389/ffgc.2019.00022 (2019).

39 Feintrenie, L., Chong, W. K. & Levang, P. Why do Farmers Prefer Oil Palm? Lessons Learnt from Bungo District, Indonesia. Small-scale Forestry 9, 379–396, doi:10.1007/s11842-010-9122-2 (2010).

40 Fong, Y. C., Khin, A. A. & Lim, C. S. Determinants of Natural Rubber Price Instability for Four Major Producing Countries. Social Sciences and Humanities 28, 1179–1197 (2020).

41 Soratana, K., Rasutis, D., Azarabadi, H., Eranki, P. L. & Landis, A. E. Guayule as an alternative source of natural rubber: A comparative life cycle assessment with Hevea and synthetic rubber. Journal of Cleaner Production 159, 271–280, doi:10.1016/j.jclepro.2017.05.070 (2017).

42 Wang, J. et al. Large Chinese land carbon sink estimated from atmospheric carbon dioxide data. Nature 586, 720–723, doi:10.1038/s41586-020-2849-9 (2020).

43 Wigboldus, S. et al. Scaling green rubber cultivation in Southwest China-An integrative analysis of stakeholder perspectives. Sci Total Environ 580, 1475–1482, doi:10.1016/j.scitotenv.2016.12.126 (2017).

44 Xu, J., Grumbine, R. E. & Beckschäfer, P. Landscape transformation through the use of ecological and socioeconomic indicators in Xishuangbanna, Southwest China, Mekong Region. Ecological Indicators 36, 749–756, doi:10.1016/j.ecolind.2012.08.023 (2014).

45 FSC. FSC Webinar on Unlocking Sustainable Natural Rubber: How to scale early successes for 2022. https://www.youtube.com/watch?v=o-tnPmnR1e4 (2021).

46 European Union. Feasibility study on options to step up EU action against deforestation. (2018).

## Methods references

1 Contains modified Copernicus Sentinel data [2020-2022] for Sentinel data. (2015).

2 Gorelick, N. et al. Google Earth Engine: Planetary-scale geospatial analysis for everyone. Remote Sensing of Environment 202, 18–27, doi:10.1016/j.rse.2017.06.031 (2017).

3 Louis, J. et al. in Proceedings living planet symposium 2016. 1–8 (Spacebooks Online).

4 Main-Knorn, M. et al. in Image and Signal Processing for Remote Sensing XXIII. 37–48 (SPIE).

5 Louis, J. et al. in IGARSS 2019-2019 IEEE International Geoscience and Remote Sensing Symposium. 8522–8525 (IEEE).

6 Xiao, C., Li, P., Feng, Z., Liu, Y. & Zhang, X. Sentinel-2 red-edge spectral indices (RESI) suitability for mapping rubber boom in Luang Namtha Province, northern Lao PDR. International Journal of Applied Earth Observation and Geoinformation 93, 102176 (2020).

7 Monthly precipitation in mm at 1 km resolution based on SM2RAIN-ASCAT 2007-2018 and IMERG 2014-2018 10.5281/zenodo.1435912.

8 Alchemi, P. & Jamin, S. in IOP Conference Series: Earth and Environmental Science. 012030 (IOP Publishing).

9 Niu, F., Röll, A., Meijide, A. & Hölscher, D. Rubber tree transpiration in the lowlands of Sumatra. Ecohydrology 10, e1882 (2017).

10 Zanaga, D. V. D. K., Ruben; De Keersmaecker, Wanda; Souverijns, Niels; Brockmann, Carsten; Quast, Ralf; Wevers, Jan; Grosu, Alex; Paccini, Audrey; Vergnaud, Sylvain; Cartus, Oliver; Santoro, Maurizio; Fritz, Steffen; Georgieva, Ivelina; Lesiv, Myroslava; Carter, Sarah; Herold, Martin; Li, Linlin; Tsendbazar, Nandin-Erdene; Ramoino, Fabrizio; Arino, Olivier. ESA WorldCover 10 m 2020 v100. (doi:10.5281/zenodo.5571936). (2021).

11 Lan, G. et al. Main drivers of plant diversity patterns of rubber plantations in the Greater Mekong Subregion. Biogeosciences 19, 1995–2005 (2022).

12 Yang, J., Xu, J. & Zhai, D.-L. Integrating phenological and geographical information with artificial intelligence algorithm to map rubber plantations in Xishuangbanna. Remote Sensing 13, 2793 (2021).

13 Bey, A. et al. Collect earth: Land use and land cover assessment through augmented visual interpretation. Remote Sensing 8, 807 (2016).

14 Markert, K. N. et al. in AGU Fall Meeting Abstracts. IN23C–0099.

15 Saah, D. et al. Collect Earth: An online tool for systematic reference data collection in land cover and use applications. Environmental Modelling & Software 118, 166–171 (2019).

16 Yu, L. & Gong, P. Google Earth as a virtual globe tool for Earth science applications at the global scale: progress and perspectives. International Journal of Remote Sensing 33, 3966–3986 (2012).

17 Team, P. Planet Application Program Interface: In Space for Life on Earth. San Francisco, CA. https://api.planet.com. (2017).

18 Landsat-5-7-8 image courtesy of the U.S. Geological Survey.

19 Turubanova, S., Potapov, P. V., Tyukavina, A. & Hansen, M. C. Ongoing primary forest loss in Brazil, Democratic Republic of the Congo, and Indonesia. Environmental Research Letters 13, 074028 (2018).

20 Hansen, M. C. et al. High-resolution global maps of 21st-century forest cover change. science 342, 850–853 (2013).

21 Kennedy, R. E., Yang, Z. & Cohen, W. B. Detecting trends in forest disturbance and recovery using yearly Landsat time series: 1. LandTrendr—Temporal segmentation algorithms. Remote Sensing of Environment 114, 2897–2910 (2010).

22 Kennedy, R. E. et al. Implementation of the LandTrendr algorithm on google earth engine. Remote Sensing 10, 691 (2018).

23 Zhu, Z., Wang, S. & Woodcock, C. E. Improvement and expansion of the Fmask algorithm: Cloud, cloud shadow, and snow detection for Landsats 4–7, 8, and Sentinel 2 images. Remote sensing of Environment 159, 269–277 (2015).

24 BirdLife International (2022) World Database of Key Biodiversity Areas. Developed by the KBA Partnership: BirdLife International, International Union for the Conservation of Nature, American Bird Conservancy, Amphibian Survival Alliance, Conservation International, Critical Ecosystem Partnership Fund, Global Environment Facility, Re:wild, NatureServe, Rainforest Trust, Royal Society for the Protection of Birds, Wildlife Conservation Society and World Wildlife Fund. March 2022 version. Available at http://keybiodiversityareas.org/kba-data/request. (2022).

