## Supplementary Information for "New high-resolution maps show that rubber causes significant deforestation"

##### Extended Data Note | Definitions of 'forest' and 'deforestation'

The definition of forest is a critical issue, for example when comparing different deforestation estimates (e.g., in Table 2). It is very difficult to distinguish plantation forest and agricultural tree crops from natural forest and studies deal with this in different ways. In addition, studies use different baseline tree cover thresholds.

On the first point, Goldman et al. (2020)[1] opted to use the term 'deforestation' to mean all tree cover loss, including rotational tree crop and plantation clearing. Hurni & Fox (2018)[2] on the other hand track changes in a range of tree crops and thereby minimise their confounding influence. Pendrill et al. (2019)[3] took steps to exclude tree plantations in Indonesia and Malaysia, and elsewhere use 'deforestation' to mean all tree cover loss. As described in the Methods, in this study we minimise the erroneous inclusion of plantations by (1) tracking only the first deforestation date going back as far as the Landsat imagery allows (1988), and (2) only counting pixels above an NBR threshold (prior to the change detection) to reduce the inclusion of plantations for which deforestation occurred prior to 1988.

Our deforestation estimates for Indonesia and Malaysia are higher than those of Pendrill et al. (2019)[3]. However, even if one assumed that for these two countries our data presents an overestimate (i.e. includes plantation rotation), and if one replaced our estimates for Indonesia and Malaysia with those of Pendrill et al. (2019)[3], our overall figure for annual rubber-related deforestation would still be more than twice as high. Furthermore, even if one assumed that no deforestation took place at all in Indonesia, Malaysia and also Thailand, our annual deforestation figure would still be marginally above the total figure provided by Pendrill et al. (2019)[3]. It is also noteworthy that although Hurni and Fox (2018)[2] employ a strict approach for dealing with tree crops, their deforestation figures are generally in line, and in some areas (Cambodia) above, ours.

On the second point: Pendrill et al. (2019)[3] use a stricter definition of baseline tree cover than employed in this study (using a higher canopy cover threshold of 25% versus 10% in this study); however, according to a sensitivity analysis conducted by the authors themselves a lower (10%) tree cover threshold would generally not lead to big differences in estimated deforestation (apart from in Africa where the difference is more notable)[3]. Goldman et al. (2020)[1] use an even stricter tree cover threshold of 30%, but their deforestation figures are generally consistent with (or above) our figures; this may be because the effect of a stricter tree cover threshold is balanced by a less strict approach to plantation inclusion (Goldman et al. (2020) use a baseline of 2000 as opposed to the baseline of 1988 used in this study).

**Not yet peer-reviewed and potentially still subject to corrections**

#### References


1. Goldman, E., Weisse, M. J., Harris, N. & Schenider, M. Estimating the Role of Seven Commodities in Agriculture-Linked Deforestation: Oil Palm, Soy, Cattle, Wood Fiber, Cocoa, Coffee, and Rubber. (Technical Note. Washington, DC: World Resources Institute., 2020).
2. Hurni, K. & Fox, J. The expansion of tree-based boom crops in mainland Southeast Asia: 2001 to 2014. *Journal of Land Use Science* **13**, 198-219, doi:10.1080/1747423x.2018.1499830 (2018).
3. Pendrill, F., Persson, U. M., Godar, J. & Kastner, T. Deforestation displaced: trade in forest-risk commodities and the prospects for a global forest transition. *Environmental Research Letters* **14**, doi:10.1088/1748-9326/ab0d41 (2019).


**Not yet peer-reviewed and potentially still subject to corrections**

### Extended Data tables

**Extended Data Table 1 | Confusion matrix for mapping rubber across Southeast Asia.**

| | | Reference samples | | | | |
|---|---|---|---|---|---|---|
| | | Forest | Rubber | Others | Sum | User's accuracy |
| Map | Forest | 338 | 19 | 0 | 357 | 0.9468 |
| | Rubber | 2 | 302 | 0 | 304 | 0.9934 |
| | Other | 5 | 41 | 0 | 46 | NA |
| | Sum | 345 | 362 | 0 | 707 | |
| Producer's accuracy | | 0.9797 | 0.8343 | NA | | Overall accuracy: 0.9052 |

**Extended Data Table 2 | Confusion matrix for mapping deforestation associated with rubber across Southeast Asia.**

| | | Reference samples | | | |
|---|---|---|---|---|---|
| | | Deforestation ≤ 2000 | Deforestation > 2001 | Sum | User's accuracy |
| Map | Deforestation ≤ 2000 | 509 | 15 | 524 | 0.9714 |
| | Deforestation > 2001 | 139 | 41 | 180 | 0.2278 |
| | Sum | 648 | 56 | 704 | |
| Producer's accuracy | | 0.7855 | 0.7321 | | Overall accuracy: 0.7813 |

**Extended Data Table 3 | Area of rubber-related deforestation for individual countries in Southeast Asia.** The 95% confidence Interval (CI) was calculated using the accuracy estimates presented in Extended Data Table 2.

| **Country** | **Deforestation 2001-2016 (ha, A)** | **Deforestation / Rubber** | **Deforestation in KBA (ha, C)** | **C/A** |
|---|---|---|---|---|
| China | 169,294 | 15% | 16,055 | 9% |
| Myanmar | 118,588 | 15% | 14,286 | 12% |
| Cambodia | 244,444 | 40% | 31,482 | 13% |
| Viet Nam | 218,674 | 14% | 9,039 | 4% |
| Laos | 136,137 | 24% | 9,689 | 7% |
| Malaysia | 316,739 | 32% | 15,760 | 5% |
| Thailand | 549,964 | 15% | 29,559 | 5% |
| Indonesia | 1,014,147 | 20% | 91,592 | 9% |
| **Southeast Asia (95% confidence interval)** | **2,767,986 ± 239,189** | **19.04 ± 11.81%** | **217,461** | **8 ± 1%** |

Not yet peer-reviewed and potentially still subject to corrections

#### Extended Data figures

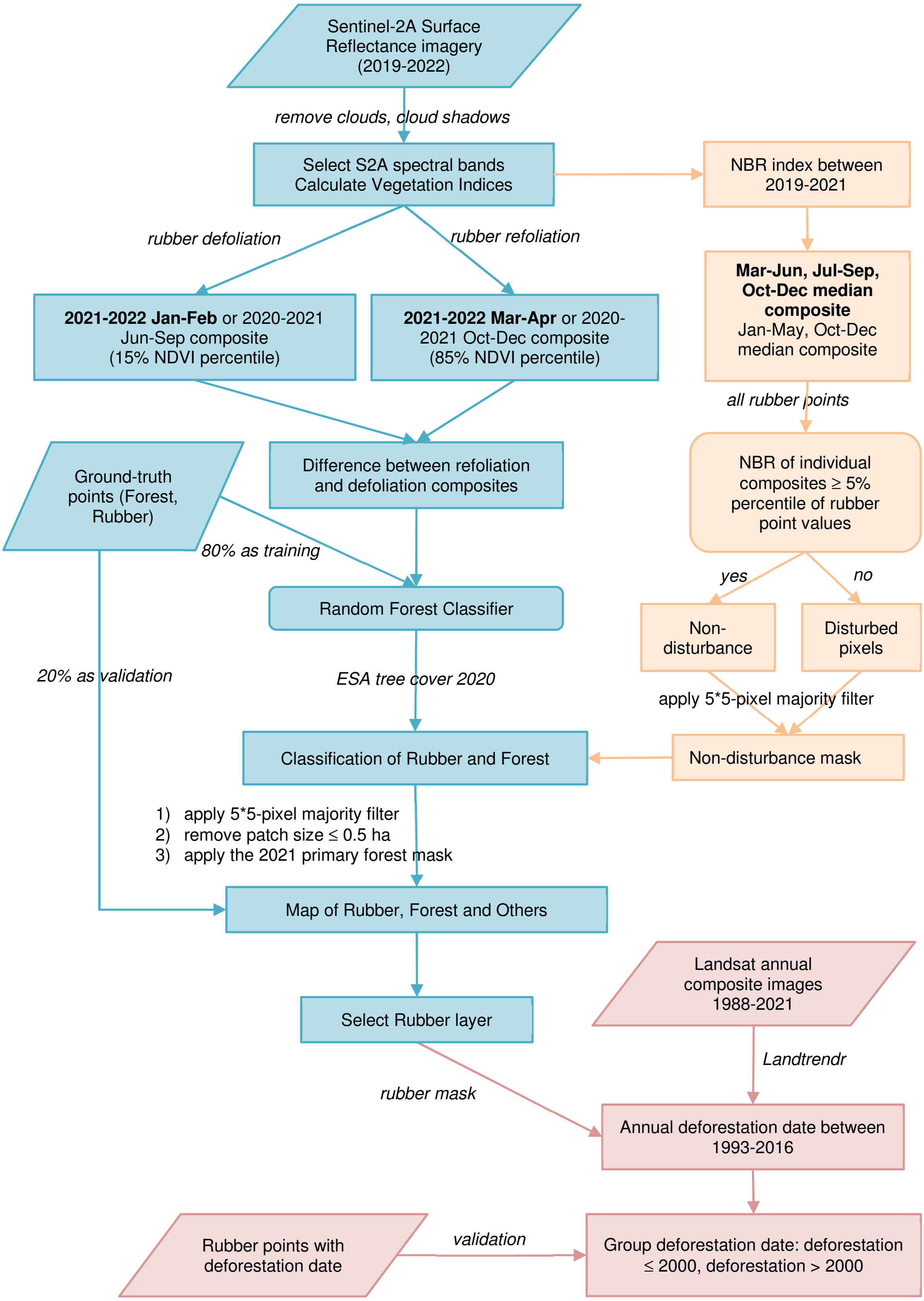


**Extended Data Fig.1 | Methodology flow for mapping rubber (blue), generating non-disturbance mask (green) and estimating deforestation (orange).** Different image composites were used for region-A (defoliation between January-February) and region-B (defoliation between June-September). All processing was done in Google Earth Engine.

**Not yet peer-reviewed and potentially still subject to corrections**

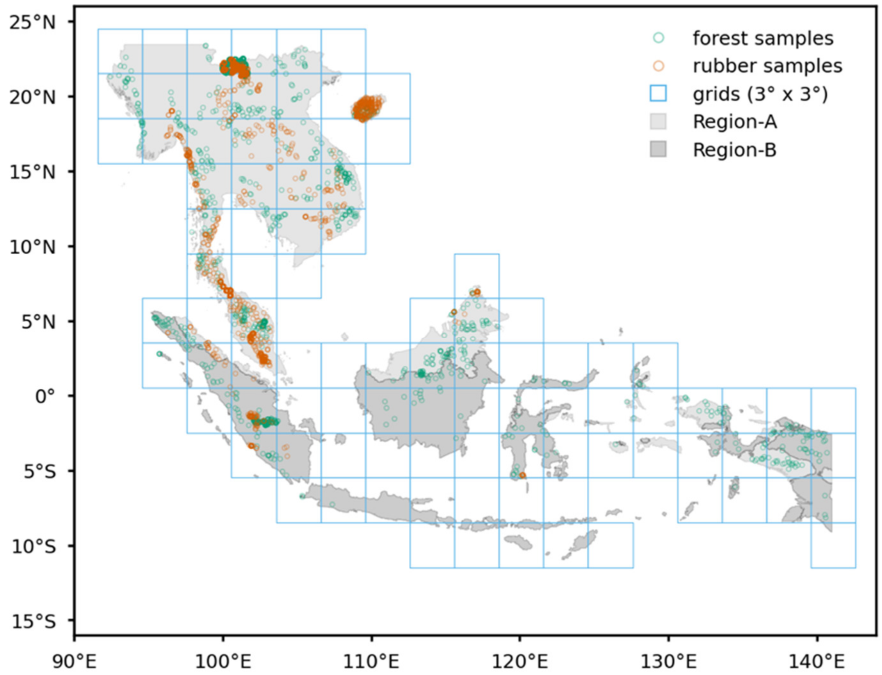


**Extended Data Fig.2 | Rubber phenology regions, grids, and sampling points.** As rubber phenology varies across Southeast Asia we divided the study area into two regions using OpenLandMap Monthly Precipitation[7]. Region-A: rubber defoliation was assumed to occur between January-February and refoliation between March-April. Region-B: rubber defoliation was assumed to occur between June-September and refoliation between October-December. The algorithm was run separately for 3 by 3-degree grid cells (in blue). The forest and rubber sample ground-truth points were used for training the algorithm (80%) and subsequently validating the map (20%).

Not yet peer-reviewed and potentially still subject to corrections
